# An invariant relationship between NonREM and REM sleep and the wave model of their dynamics

**DOI:** 10.1101/2023.01.19.524817

**Authors:** Vasili Kharchenko, Irina V. Zhdanova

## Abstract

Explaining the complex structure and dynamics of sleep, which consist of alternating and physiologically distinct NonREM and REM sleep episodes, has posed a significant challenge. In this study, we demonstrate that a single wave model concept captures the distinctly different overnight dynamics of the four primary sleep measures - the duration and intensity of NonREM and REM sleep episodes - with high quantitative precision. Additionally, the model accurately predicts how these measures respond to sleep deprivation or abundance. Furthermore, the model passes the ultimate test, as its prediction leads to a novel experimental finding—an invariant relationship between the duration of NonREM episodes and the intensity of REM episodes, the product of which remains constant over consecutive sleep cycles. These results suggest a functional unity between NonREM and REM sleep, establishing a comprehensive and quantitative framework for understanding normal sleep and sleep disorders.

## INTRODUCTION

Despite significant progress in understanding the brain mechanisms involved in sleep regulation^1^, the physiological function and dynamics of sleep within the sleep-wake homeostasis framework^2^ remain uncertain and a topic of debate^3,4^. Sleep has a complex structure known as sleep architecture and, in humans, typically includes five to six sleep cycles per night (Fig. 1a). Each sleep cycle consists of two types of sleep: it starts with non-rapid eye movement sleep (NREMS) and followed by rapid eye movement sleep (REMS).

**Figure 1.**
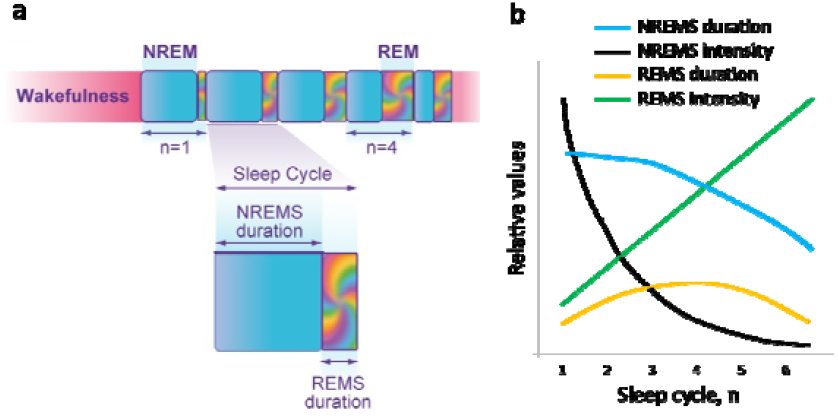
Human sleep architecture and overnight dynamics of the four primary sleep measures. a. Schematics of consolidated sleep pattern, consisting of consecutive sleep cycles, each including longer NREMS (cyan) and shorter REMS (multicolor) episodes. Five sleep cycles are shown, n. b. Schematics of typical changes in sleep measures over consecutive sleep cycles of regular nighttime sleep: NREMS duration (cyan), NREMS intensity (% of Slow Wave Sleep, black), REMS duration (orange) and REMS intensity (REM density, green).

In contrast to Wake state, both NREMS and REMS are characterized by low perception of the environment, but they are otherwise remarkably different states of the organism. Gradual disengagement from the environment in NREMS is associated with slow brain activity and reduced muscle tone, though ability for motor activity is preserved. REMS is known as paradoxical sleep since manifests as high brain activity and rapid eye movements, active dream mentation, irregular heart rate and respiration, and sexual arousal, all on the background of further reduction of perception, loss of muscle tone and thermoregulation^5,6^. This unique simultaneous presence of wake-like and sleep-like features makes the nature and significance of REMS particularly puzzling.

Understanding the intricate patterns of sleep dynamics represents a major challenge in the field. For several decades, there has been a recognized need for a comprehensive mathematical model that can describe both the Sleep-Wake and NREMS-REMS state transitions. Such unifying model could provide a framework for elucidating the underlying mechanisms and help identify key factors and variables that affect the timing, duration, and quality of the Sleep state and its components.

The major advance was achieved by the two-process model of sleep regulation^2^, which is based on the empirical analysis of the intensity of NREM sleep. It proposed that, during wakefulness, the homeostatic mechanisms define exponential rise in “sleep pressure” or “need for sleep”. This process was also described as the rise in “Wake state instability”^7^, manifesting as increase in sleepiness, performance deficits and variability in motor and cognitive responses^8-10^. As sleep deprivation continues, subjective and objective symptoms of state instability diversify, involving the autonomic regulation, microvascular and cardiovascular functions, and more^11-13^. The most prominent indicators of high Wake state instability are the microsleep intrusions and involuntary sleep initiations that increase in frequency and duration over the extended wakefulness^14^. Considering that the main driver for any state transition, whether in physics, chemistry or biology, is the relative instability of one state compared to the other, these experimental observations suggest that the accumulation of Wake state instability facilitates the spontaneous transition to a more stable Sleep state.

These important models and concepts, however, did not incorporate REM sleep and, over the years, other theoretical approaches have been employed to address the complexities of the NREMS – REMS dynamics^2,15-18^. Those considered the potential homeostatic relationship between REMS and NREMS or inter-REMS interval^19^, postulation of a specialized REMS oscillator that is active only during sleep^20^, and modeling of ultradian oscillations based on classical mechanics of a particle oscillating between three potential wells representing Wake, NREMS and REMS^16^. Although these models can simulate realistic patterns of sleep-like oscillations, they have not been able to quantitatively reproduce the typical sleep architecture or functionally unite NREMS and REMS.

In our approach to sleep dynamics, we relied on the earlier concepts of homeostatic regulation of sleep pressure and state instability^2,7^. We then suggested that quasi-periodic alternations between the two types of sleep may reflect the wave dynamics of the global physiological states, Sleep and Wake. These global states are known to involve numerous interdependent negative feedback loops that define complex oscillations of critical biological variables around equilibrium setpoints on various levels, from molecular to systemic. To maintain the dynamic stability of an organism within physiological limits, it is essential for these homeostatic loops to be in coherence^21,22^. It is well established in physics that the coherent behavior of multiple oscillators typically leads to the formation of a wave and thus global physiological states may present wave behavior.

Wave processes are prevalent in biological systems, facilitating rapid long-range spatiotemporal coordination and signal preservation^23^. Since both Sleep and Wake states are stochastic processes, predicting the precise timing and duration of overall sleep or its components is challenging. However, on average, normal overnight sleep exhibits typical changes in the duration and intensity of NREM and REM sleep episodes (see Fig. 1b). This pattern resembles the behavior of probability waves^24^. This would not be surprising, considering that both experimental and theoretical physics have shown that stochastic classical systems can mimic probability wave dynamics and be accurately described using wave mechanics^25-29^.

Developing a comprehensive and unifying wave model of sleep dynamics requires incorporating the interactions between Sleep and Wake states, as well as between NREM and REM sleep. Such model must then undergo rigorous quantitative testing of its predictions against the dynamics of multiple independent experimental measures representing both NREM and REM sleep. Notably, the dynamics of the four primary sleep measures are strikingly distinct from one another (see Fig. 1b). Over successive sleep cycles, there is a gradual decline in NREM episode duration, accompanied by a rapid exponential decrease in NREM intensity, a linear increase in REM intensity, and a bell-shaped pattern of REM episode duration.

Here we show that the wave model of sleep dynamics can accurately and quantitatively describe typical human sleep architecture and dynamics of all four primary sleep measures. It then accurately predicts the effects of sleep deprivation and sleep abundance on these sleep measures. Importantly, the model suggests a functional link between NREMS and REMS, predicting an invariant relationship between NREMS duration and REMS intensity over the course of the night, which we now demonstrate experimentally.

## RESULTS

### Modeling Sleep-Wake homeostasis through interacting waves

We modeled the dynamics of Wake and Sleep states as interaction of two waves, each shaped by the Morse potential well, a mathematical function widely used in physics and chemistry to describe atomic interactions in molecules near their equilibrium. Figure 2 illustrates that low values of *x*, the regulating parameter of state stability, promote the Wake state, while high values favor the Sleep state (Methods 1-3). During daytime wakefulness, the collective action of homeostatic, circadian, and environmental forces drives *x* away from the parametric region of stable Wake state. The exponential rise in Wake state instability is then formally described as an increase in the model parameter or, in analogy to physical systems, the energy of state instability accumulated due to the work of the driving forces.

**Figure 2.**
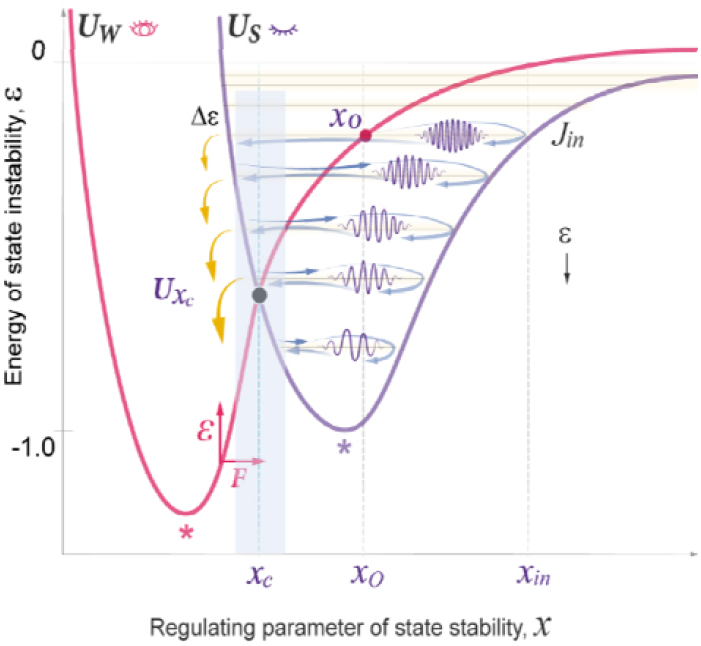
The wave model of sleep dynamics. In the Wake state (*U*_*W*_, red potential), the driving force (horizontal red arrow) increases the value of the regulating parameter *x* (horizontal axis) beyond the Sleep-Wake homeostatic equilibrium setpoint *x*_*c*_ (vertical cyan line) and the *x*_*c*_ region of efficient interaction of two states (gray area). This rises the energy of state instability (vertical red arrow and vertical axis) above the homeostatic energy threshold (*Ux*_*c*_, black dot; the crossing poin of *U*_*W*_ and *U*_*S*_ curves). Sleep (*U*_*S*_, purple potential) is initiated at *x*_*o*_ (red dot) of the initial energy level *J*_*in*_ (first sleep cycle). Relaxation of *x* and occurs in the form of sleep wavepacket propagating along energy levels (blue arrows), where *x*_*in*_ – maximal deviation. When the wavepacket reaches *x*_*c*_, a portion of its energy (yellow arrows) is released. In the Morse potential, energy gaps between levels increase linearly over the course of energy relaxation (downwards black arrow) and correlate negatively with the width of the potential well. Stars– maximal stability of the corresponding state. The schematics serves to only illustrate the concept. For actual position of *Ux*_*c*_ and *J*_*in*_ within *U*_*S*_ see Supplemental figure 1.

At night, the driving forces subside, and the energy accumulated during wakefulness is spontaneously transferred from the metastable Wake to the more stable Sleep state at _*o*_ of the initial level *j*_*in*_, generating a sleep wavepacket with energy (Fig. 2; Methods 4, 5). This initiates energy relaxation through a series of quasi-periodic oscillations, the sleep cycles, with each cycle resulting in a transition to a lower energy level. Eventually, the relaxation of the accumulated energy during Sleep can move the system back to the original state of stable Wake state, closing the homeostatic Sleep-Wake loop.

Figure 3a illustrates the two simple model parameters - the width of the potential well (and the number of the top occupied energy level (*j*_*in*,_) –that are sufficient to define the character of oscillations within the Morse potential, including the period, amplitude and the energy lost during the transition to a lower level (energy gap). The model predicted that these two parameters depend on the two principal modulators of sleep dynamics – the habitual sleep duration and the duration of wakefulness prior to the experimental sleep night. Thus, once the analysis of the dynamics of one primary sleep measure would define the values for these two model parameters for a given experimental group, they allow to predict the dynamics of other primary sleep parameters of the same dataset.

**Figure 3.**
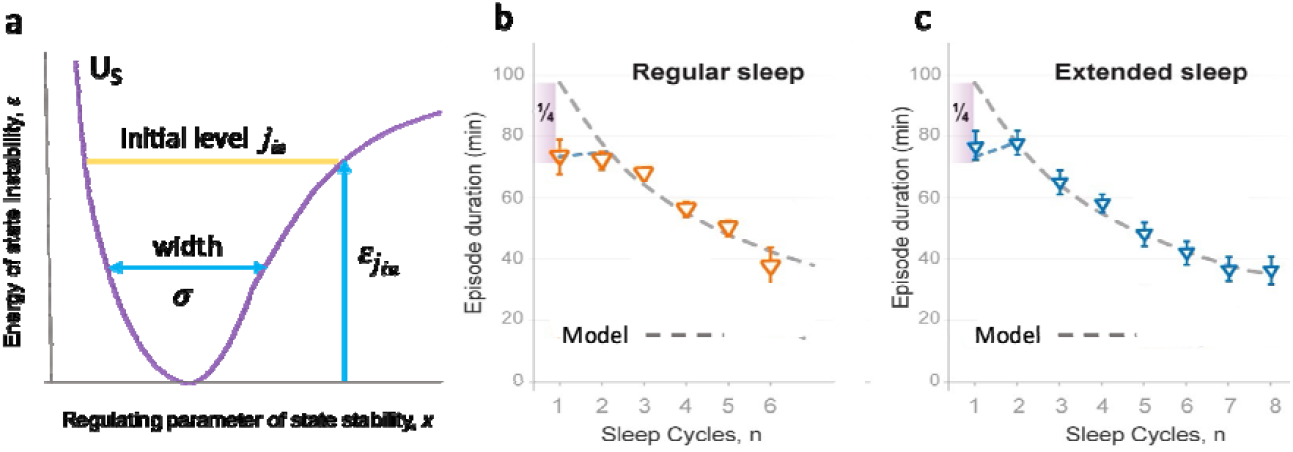
The wave model provides accurate quantitative description of the dynamics in NREMS episode duration. a. Two model parameters that allow to describe NREMS episode duration: – width of *U*_*S*_ potential (cyan doubl - arrow); - energy of state instability reached by sleep initiation (vertical cyan arrow). *j*_in_ – number of initial energy level at which sleep is initiation (yellow line). b. Regular sleep: Theoretical curve (dashed line) and experimental data (min, mean SEM) for NREMS episode durations (triangles), as a function of the sleep cycle order number n (horizontal axis). Due to *j*_in_ being populated at a variable point _o_ (see Fig. 2), the first experimental NREMS episode duration is, on average, ¼-than shorter than the theoretical one for this energy level. Data collected in 24 young healthy subjects, 39 nine-hour nights (Method 16). R^2^ value: 0.905. c. Extended sleep: Theoretical curve (dashed line) and experimental data (min, mean SEM) for NREMS episode durations (triangles), as a function of the sleep cycle order number n (horizontal axis). Data collected in 11 young healthy subjects over 308 fourteen-hour nights, as reported by Barbato & Wher^30^. First NREMS episode duration is explained in b. R^2^ value: 0.993.

### The wave model accurately describes the dynamics of NREMS episode duration

To test the model against the physiological sleep measures, we used group data, as the stochastic nature of individual sleep patterns requires additional considerations, such as probabilistic treatment of model parameters and the influence of intrinsic and environmental perturbations or noise. First, we tested the model against the dynamics of NREMS episode duration in our group of young healthy volunteers with normal sleep of regular duration (Fig. 3b, Method 16).

The decrease of NREMS episode duration over consecutive cycles (Fig. 1b) is similar to the decline in the period of oscillations of the sleep wavepacket along the regulating parameter *x* in the Morse potential (Supplemental Fig. 2). As predicted, the right combination of the two model parameters *σ* and *j*_*in*_ (Fig. 3a), allowed for the accurate depiction of relative changes in NREMS episode durations of this experimental dataset (Fig. 3b; χ^2^ goodness of fit test P>0.88, first episode excluded). As expected, the duration of the first NREMS episode was curtailed by approximately a quarter of the theoretically predicted whole period due to the position of the sleep onset (*x*_*o*_) in between the two potential walls (Fig. 2).

To validate the model against independent observations and in subjects with different habitual sleep duration and prior wakefulness, we then tested the model against the dynamics of NREMS episode duration in a dataset collected by Barbato and Wehr^30,31^. In this experimental group, subjects displayed extended sleep duration over four weeks of daily exposure to 14-hour sleep-favoring conditions. The model predictions were again affirmed, providing excellent fit to the experimental data (Fig. 3c), with high statistical significance for this experimental group (χ^2^ goodness of fit test P>0.99, first episode excluded) reflecting the high statistical power of this outstanding dataset (308 sleep nights). The model prediction that the first NREMS episode is, on average, curtailed by approximately a quarter of that predicted for the initial energy level was also confirmed in this analysis. Moreover, as predicted, longer sleep duration and more sleep cycles in the extended sleep group were associated with increased and *j*_*in*_ (Supplemental Table 1).

Collectively, our findings demonstrated that the overnight decline in the duration of NREMS episodes can be accurately and quantitatively described by the consecutive periods of the wavepacket oscillations within the Morse potential based on the right combination of the two model parameters, *σ* and *j*_*in*_. However, for any model to claim adequate representation of an overall process, it is necessary to predict the dynamics of several independent measures characterizing this process and that are not part of the development of the model. Hence, to demonstrate that the two interacting probability waves can indeed describe the overall sleep architecture, the model had to precisely predict all four primary sleep measures. To date, no sleep model has attained this validation threshold.

### Quantitative predictions of the dynamics of NREMS intensity

The intensity of a wave is directly proportional to the square of its amplitude L^2^, as per wave mechanics. By already knowing the two model parameters *σ* and *j*_*in*_ that characterizes both datasets, those for regular and extended sleep (Fig. 3), we calculated the amplitude of the wavepacket oscillations (Fig. 4a; Method 13). The model also predicted that the initial wave intensity and the rate of its decline positively correlate with the maximal energy reached at the initial level, 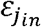. We then hypothesized that the intensity of the wavepacket oscillation at each energy level corresponds to NREMS intensity and expressed it as κL^2^, where the coefficient κ is inversely proportional to |*ε*|, the absolute value of wavepacket energy (Method 13). Note that, in general, *ε* values in a potential well are presented as negative, so an increase in *ε* leads to a lower |*ε*| (Fig. 2).

**Figure 4.**
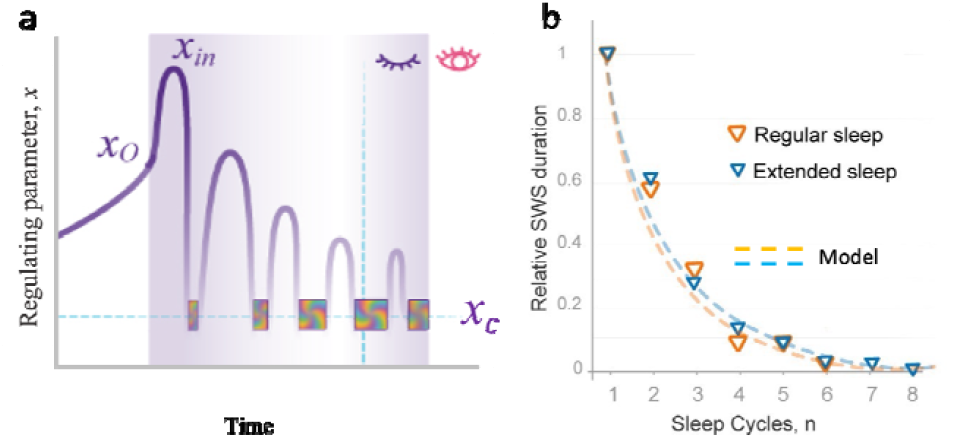
The wave model of sleep provides accurate quantitative prediction of the dynamics of NREMS episode intensity. a. Schematics of the time-dependent changes in the amplitude (L) of the regulating parameter of state stability, *x* (vertical axis) during Sleep (purple area). *x*_o_ – sleep onset, *x*_*in*_ *–* amplitude of the first cycle, *x*_c_ – equilibrium region of homeostatic threshold (horizontal dashed line), vertical dashed line - time of *Ux*_c_ passage (see Fig. 2), multicolor blocks – REMS episodes. b. The decline in NREMS intensity is proportional to L^2^. Theoretical curves (dashed lines) and experimental data (triangles) for the decline in NREMS intensity (proportional to L^2^) over consecutive sleep cycles (*n*; horizontal axis) of regular sleep (orange, as in Fig. 3b) and extended sleep (blue; as in Fig. 3c; Barbato et al.^31^). Mean group data for Slow Wave Sleep duration is normalized to the first sleep cycle. R^2^ values: Regular sleep: 0.973; Extended sleep: 0.990.

Conventionally, NREMS intensity is evaluated based on the power of slow-wave activity (SWA) in the brain cortex or the duration of slow-wave sleep (SWS)^32^. To test our quantitative prediction regarding NREMS intensity, we compared the SWS durations per cycle in the experimental groups with regular and extended sleep to our model’s predictions. Within each group, NREMS intensity was normalized to the first sleep cycle. We found that our theoretical curves were in good agreement with experimentally observed dynamics of NREMS intensity for both experimental groups (Fig. 4b, P>0.99 for both datasets).

Together, accurate description of the dynamics of NREMS episode duration by a combination of two model parameters generated precise prediction of the dynamics of NREMS intensity over consecutive sleep cycles. This also provided mathematical and conceptual explanation for the expedient decline of NREMS intensity, which flattens out mid-sleep, substantially outpacing the decline in NREMS episode duration or the overall sleep duration.

### Quantitative predictions of the dynamics of REMS intensity

Validation of the model prediction that propagation of the wavepacket at each energy level corresponds to NREMS, as per both NREMS duration and intensity (Fig. 3, 4), suggested that the position of REMS episodes in between NREMS episodes may correspond to the transition of the wavepacket from higher to lower energy level. Accordingly, an intensity of REMS was then predicted to be proportional to the magnitude of the energy released during each transition (Fig. 2). This corresponds to the energy gap between adjacent levels that depends on the same two model parameters and *j*_*in*_ (Fig. 3a). Notably, in the Morse potential, as in other semi-classical potentials, is inversely proportional to the period of oscillation and increases linearly towards lower energy levels.

Typically, REMS intensity is evaluated based on the number of eye movements per minute of REMS episode, referred to as REM density^33-36^. Since energy gaps in the Morse potential are small at high energy levels and increase linearly as the energy is relaxed (Fig. 2, 5a), the model predicted that REMS intensity should also be low at the sleep start and increase linearly over consecutive REMS episodes. This is consistent with experimental data accumulated over the past decades that shows linear increase in quantitatively assessed REMS intensity over consecutive cycles of normal sleep^33-36^ (Fig. 5b). This increase depends on sleep homeostasis, not the circadian regulation^36^.

**Figure 5.**
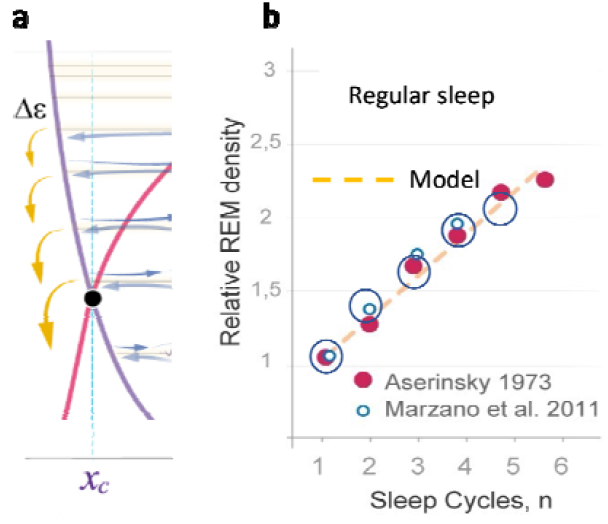
The wave model accurately predicts the slope of linear increase in REMS intensity. a. The schematics of the region of homeostatic equilibrium between Sleep and Wake states (gray *x*_*c*_ region in Fig. 3) depicting linear increase in energy release that occurs in REMS due to linear increase in energy gaps between levels of the Morse potential. b. The rate of linear increase in REMS intensity (REM density) over consecutive sleep cycles documented in three similar but independent groups, as compared to theoretically predicted for a similar group by the wave model (orange dashed line). Group 1 (large open circle, n=11 nights; R^2^ 0.985; Method 16), group 2, in Aserinsky^34^ (red circle, n=20 nights, R^2^ 0.961), group 3 in Marzano et al.^35^ (small open circle, n= 50 nights; R^2^ 0.997). Theoretical prediction was based on the increase in the energy gaps Δ*ε* over consecutive cycles, with no adjusting parameters used (n=39 nights, as in Fig. 3b). Mean group data for REM density is normalized to the first sleep cycle.

Strong validation of our model was then provided by comparing the theoretical and experimental slopes of linear increase in REMS intensity over regular sleep period (Fig. 5b). In the model, the slope is defined by the same two model parameters and *j*_*in*_ (Fig. 3a). The model also predicts that *j*_*in*_ positively correlates with the duration of prior wakefulness, while positively correlates with habitual sleep duration (Method 14). Together, this suggested that groups with similar sleep durations prior to and over the experimental period should have similar REMS intensity slopes.

To test this prediction, we compared the dynamics of quantitatively assessed REM density data for normal sleep of regular duration documented in three groups of young healthy volunteers by us (n=11 nights; Method 16), Aserinsky^34^ (n=20 nights) and Marzano et al.^35^ (n=50 nights). Figure 5b illustrates that the slope angles were almost identical between these independent experimental groups and matched well the slope predicted by the model (Method 16; P >0.99 for all datasets). The latter theoretical slope was based exclusively on the values of and *j*_*in*_ documented during the analysis of NREMS episode durations for 39 nights of regular sleep (Fig. 3b).

### The Sleep Cycle Invariant: theoretical prediction and experimental demonstration

Figure 6 illustrates the wave model prediction of the Sleep Cycle Invariant (SCI) based on the validated mathematical description of NREMS and REMS dynamics. As described above, within each sleep cycle, the NREMS episode duration corresponds to the period of oscillation of the sleep wavepacket and this period is inversely proportional to the energy gap between levels, 1/ (Method 5). On the other hand, the intensity of the subsequent REMS episode is directly proportional to the energy released at the end of the cycle, (Fig. 5). This predicts that the product of the NREMS episode duration and the intensity of the following REMS episode, should remain constant over consecutive sleep cycles and this constitutes the SCI (Fig. 6; Method 5, 14, 15).

**Figure 6.**
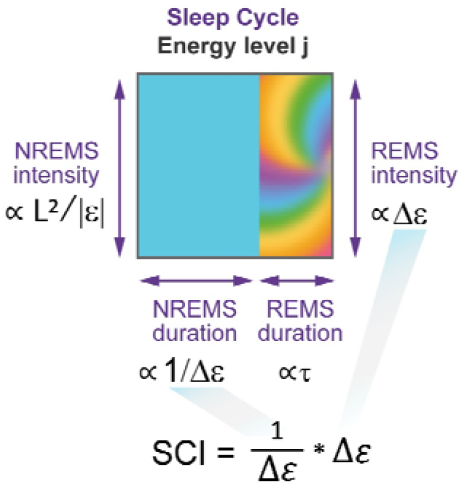
The wave model of Sleep predicts the Sleep Cycle Invariant. Experimental sleep measures (purple font) are proportional (to the wave model parameters (black font). NREMS duration is to the period of *x* oscillation and thus inverse to the energy gap, 1/. NREMS intensity is to L^2^, where L is the amplitude of *x* oscillation and is the absolute value of wavepacket energy. REMS duration is to, the lifetime of coherent superposition. REMS intensity is to energy gap,. The model predicts that the Sleep Cycle Invariant (SCI), being the product of NREMS duration and REMS intensity, should remain constant over consecutive sleep cycles. Cyan – NREMS, multicolor - REMS.

We suggested that various factors that interfere with sleep quality can modify the SCI stability and thus tested this prediction against high sleep efficiency data (not less than 93%, n=11 nights). The product of NREMS duration and REMS intensity was calculated for each sleep cycle of each individual night to evaluate the SCI.

Despite of the NREMS duration and REMS intensity showing distinct dynamics over consecutive sleep cycles, the product of these two sleep measures (SCI) remained near constant over the course of the night (Fig. 7a,b, P value > 0.92, see Statistical analysis in Methods). This confirmed the invariant relationship between NREMS and REMS over consecutive sleep cycles of normal high-efficiency sleep.

**Figure 7.**
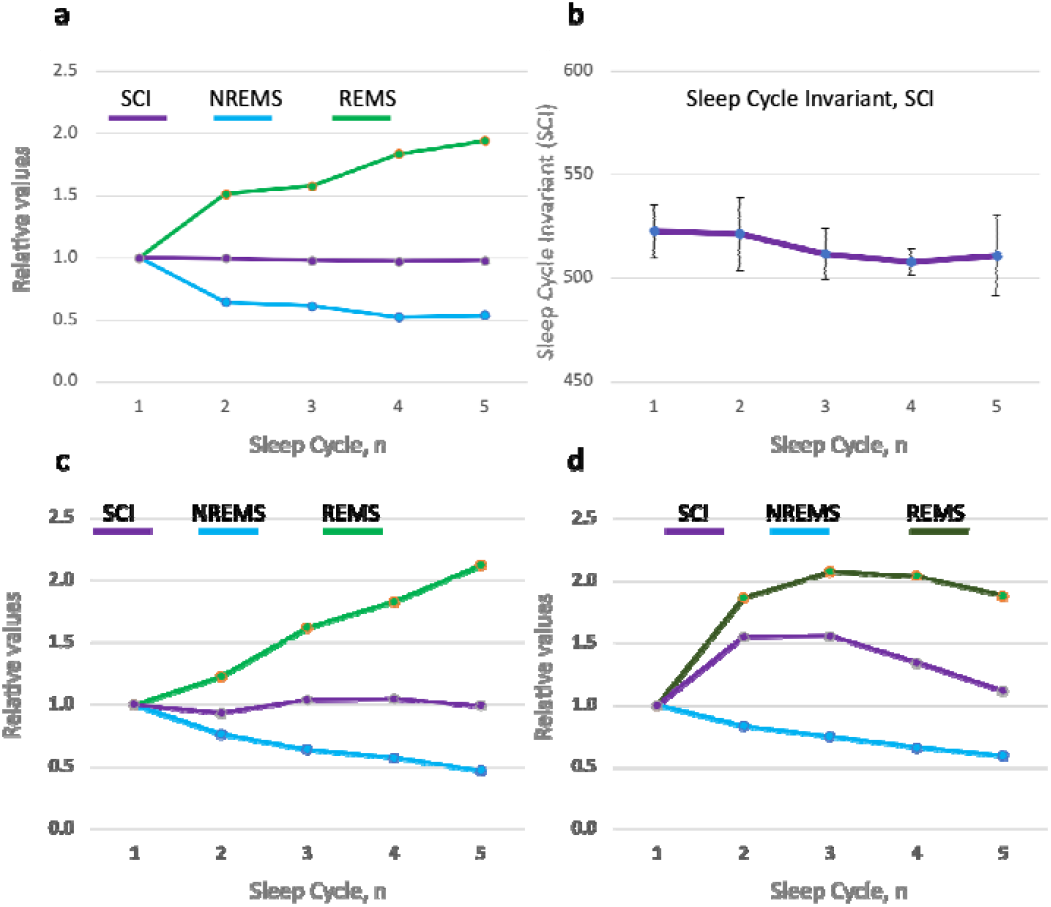
Sleep Cycle Invariant unites NREM and REM sleep. a. The product of NREMS episode duration and REMS episode intensity - the Sleep Cycle Invariant (SCI, purple), NREMS duration (cyan) and REMS intensity based on quantitative evaluation of REM density (light green) over consecutive cycles of regular sleep in a group of 11 young healthy volunteers with sleep efficiency of at least 93%; n=11 for cycles 1-4 and n=5 for cycle 5 (see Methods). Data normalized to the first sleep cycle (=1) for each parameter and presented as group means per cycle *. b. Variation in SCI over consolidated high-quality sleep. Data are presented as mean SEM; dashed line defines the mean value (515.2); the rest as in a. c. Analogous to panel A, the plot shows SCI based on NREMS duration per sleep cycle, as reported by Barbato & Wehr^*30*^ (n=308 nights; 11 young healthy volunteers), and REMS intensity per cycle, as reported by Aserinsky^*35*^ and evaluated using quantitative method (n=20 nights; 10 young healthy volunteers). d. Loss of linear time- and cycle-dependency of REMS intensity when assessed using semi-quantitative method (dark green) results in obscured NREMS-REMS invariant relationship (purple). REMS intensity was assessed in parallel with NREMS duration, as reported by Barbato et al.^*31*^ (n=208 nights; 8 young healthy volunteers). Data normalized to the first sleep cycle (=1) for each parameter and presented as group mean per cycle*. * In a-d, the duration of the first NREMS episode was multiplied by 3/4, as per the model prediction of incomplete period of the first cycle (see Methods and Fig. 2, 3).

We then aimed to evaluate SCI in previously published data on normal sleep and sleep disorders, but encountered a methodological challenge. Robust linear increase in REM density over the sleep period was originally demonstrated by Aserinsky, the discoverer of REM sleep, through quantitative assessment of rapid eye movements and was confirmed by others, also establishing its independence of the circadian phase^34-36^. However, most of the studies reported REM density based on technically more simple semi-quantitative evaluations, such as assigning a score to a range of REMs or counting number of intervals that contained REMs. These semi-quantitative measures can reveal major pathological changes in REMS intensity^37,38^ but lack sensitivity, especially at high intensity levels^39^, and thus can obscure the normal linear patterns. Consequently, we could identify only few studies that reported quantitative data on REMS intensity, and none that reported it over consecutive sleep cycles in parallel with NREMS episode durations.

We then tried a different approach to investigate the SCI in earlier studies. Since the dynamics of NREMS duration (Fig. 3a,b) and quantitatively-assessed REMS intensity (Fig. 4b) are nearly identical among groups of individuals with normal sleep, we suggested that SCI could still be observed using sleep data from different sources, provided that the groups studied were characterized by similar age and habitual sleep duration, and had normal sleep patterns. In this “hybrid” analysis, we used NREMS data from one study^30^ and REMS intensity data from another^34^, with both studies being of the highest quality and allowing subjects abundant sleep opportunity. This analysis also revealed the invariant relationship between NREMS and REMS (Fig. 7c). In contrast, when we used the results of semi-quantitative assessment of REMS intensity in combination with NREMS episode duration documented within the same large-scale study of top quality^31^, the invariant relationship was obscured (Fig. 7d). Together, these results further supported the existence of SCI in normal sleep and underscored the critical importance of quantitative assessment of REM density.

### Quantitative predictions of the dynamics of REMS duration

Finally, we have addressed the dynamics of the fourth primary sleep measure – the duration of REMS episodes. The validated model prediction that REMS intensity is directly proportional to the energy released during the transition of a wavepacket to a lower level suggested that the duration of REMS episode is determined by the duration of this energy release process.

As the wavepacket approaches the Wake state () towards the end of each NREMS episode, it enters the region of Sleep-Wake homeostatic equilibrium (_*c*_ region) where the interaction between the Sleep and Wake waves is increased (Fig. 2, 5a, 8). Such interaction typically delays the waves’ propagation (Methods 6-9), leading to their temporary coherent superposition. The Sleep-Wake interaction is strongest at the crossing of the two potentials. This can create a resonant state that further enhances the delay in the wavepacket propagation. The effect is maximal at the resonance level, the energy of which is distinct from *U*_*S*_ levels and corresponds to the homeostatic energy threshold, *U*(_*c*_) (Fig. 8; Methods 8-10). The energy dependence of the time that the wavepacket spends in the _*c*_ region of homeostatic equilibrium is then defined by the classical Lorentz resonance curve (Methods 11).

**Figure 8.**
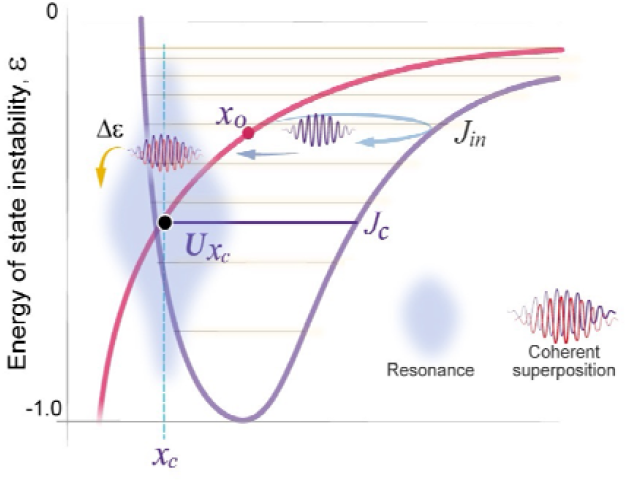
The duration of coherent superposition within the equilibrium region of Sleep and Wake states is enhanced by the resonance. Around the crossing point *U(x*_*c*_) of *U*_*S*_ (purple potential) and *U*_*W*_ (red line), strong Sleep-Wake interaction creates the resonance level and resonance state (gray area) with energy-dependent () strength (width of gray area). The propagation of the sleep-wavepacket (purple) is temporarily delayed by the resonance within the *x*_*c*_ equilibrium region. There, the wavepacket incorporates the Sleep and Wake waves and forms their coherent superposition (red-purple wavepacket). This represents REM sleep, associated with the release of a portion of energy (yellow arrow). This is a schematic representation; for actual typical position of *J*_*c*_ see Supplemental Fig. 1.

The same two model parameters, and *j*_*in*_ (Fig. 3a) characteristic of a specific dataset predicted the energy relaxation dynamics as a function of the cycle number, *n*. Fitting the right combination of the resonance energy level and the width of the resonance (Fig. 9a) then allowed for quantitative description of the non-linear dynamics of REMS episode duration (Methods 12).

**Figure 9.**
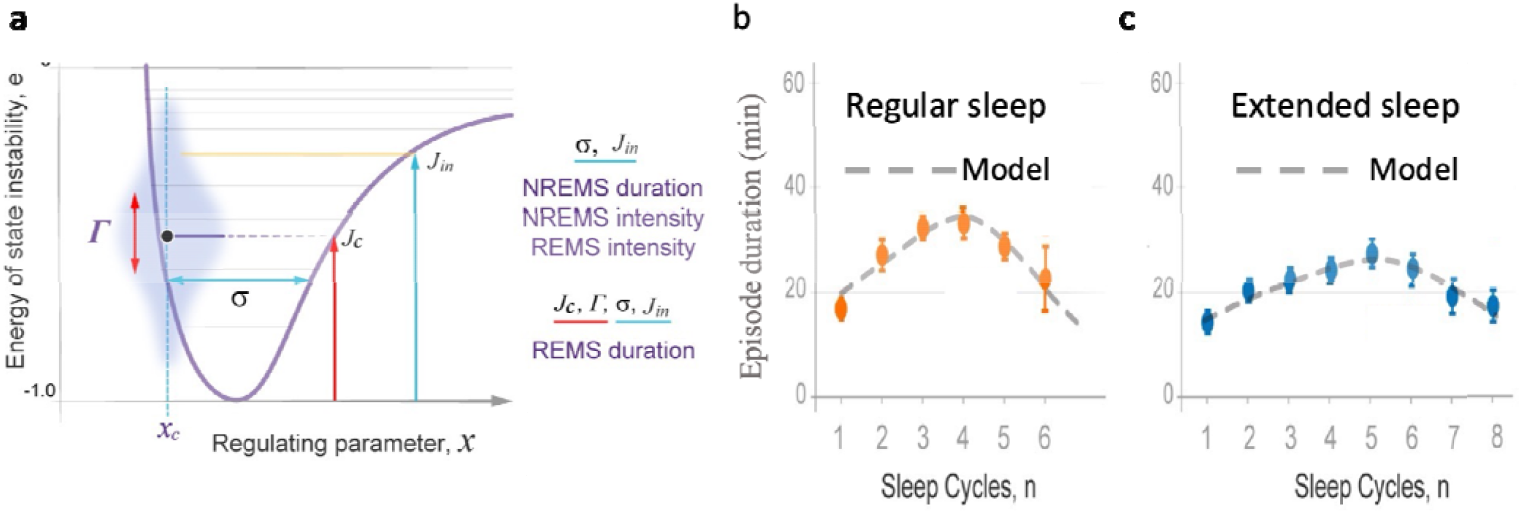
Accurate quantitative prediction of the durations of REMS episodes by the wave model of sleep dynamics. a. In addition to two model parameters, – width of *U*_*S*_ potential (cyan double-arrow) and *J*_in_ – initial energy level at which sleep is initiation (yellow line), that are required to predict the duration of NREMS episodes, intensity of NREMS and REMS (see Fig. 4-6), quantitative description of REMS episode duration requires two more model parameters: - resonanc width (red vertical double-arrow) and energy of the resonance level (red vertical arrow). Gray area – the resonance region, with wider area depicting stronger resonance. Black dot - homeostatic energy threshold *U(x*_*c*_). b. Regular sleep: Theoretical curve (dashed line) and experimental data (min, mean SEM) for REMS episode durations (circles), as a function of the sleep cycle order number n (horizontal axis). Data collected in 24 young healthy subjects, 39 nine-hour nights (Method 16). R^2^ value: 0.811. c. Extended sleep: Theoretical curve (dashed line) and experimental data (min, mean SEM) for REMS episode durations (circles), as a function of the sleep cycle order number n (horizontal axis). Data collected in 11 young healthy subjects, 308 fourteen-hour nights, as reported by Barbato and Wher^30^. R^2^ value: 0.965.

This analysis was applied to both regular and extended sleep datasets (same as in Fig. 3b,c; 4b), and the model accurately described both patterns of REMS episode duration (Fig. 9b,c; goodness of fit test P>0.95 and P>0.99, respectively). The experimentally observed bell-shaped patterns of REMS duration were consistent with the Lorentz resonance curve, according to which the closer the energy of the wavepacket is to the energy of the resonance level, the longer the presence of the wavepacket in the homeostatic equilibrium region, and thus the longer the duration of REMS episode. Accordingly, maximal REMS duration was observed around the homeostatic energy threshold, *Ux*_*c*_. Lower peak of the bell-shaped curve in extended sleep (Fig. 9c) was consistent with wider resonance in a wider potential well that has more energy levels and smaller energy gaps (Supplemental Table 1).

### The model predicts the effects of sleep deprivation or abundance on sleep architecture

Validating a model under perturbed conditions is essential for ensuring its robustness and reliability. In this report, we outline the model’s predictions for only acute perturbations under entrained conditions when sleep is initiated at habitual bedtime. These conditions provide a stable adaptive synergy between circadian and homeostatic regulation of sleep. The wave model demonstrated high accuracy in predicting the effects of acute sleep deprivation and sleep abundance on sleep architecture that have been extensively documented in various experimental settings^32-34,40,41^.

Specifically, the model predicted that the recovery sleep following sleep deprivation starts with lower REMS duration and intensity but higher NREMS intensity. Figure 10 illustrates that higher Wake state instability reached following prolonged wakefulness and thus higher energy of the initial level corresponds to weaker resonance, hence shorter initial REMS episode duration. Smaller energy gaps between higher energy levels of the Morse potential result in lower initial REMS intensity (Fig. 5). However, higher initial energy and higher amplitude of the wavepacket oscillations at higher levels lead to higher NREMS intensity (Fig. 4). The initial NREMS episode duration is a less predictable parameter following sleep deprivation mainly due to increased variation in, the position of spontaneous sleep onset (Fig. 2), or occurrence of exceptionally long NREMS episodes due to the well-known “skipped REMS episode” phenomenon. The latter can be explained by a very weak resonance at high energy levels that leads to very brief coherent superposition and thus REMS duration too short to be reliably documented.

**Figure 10.**
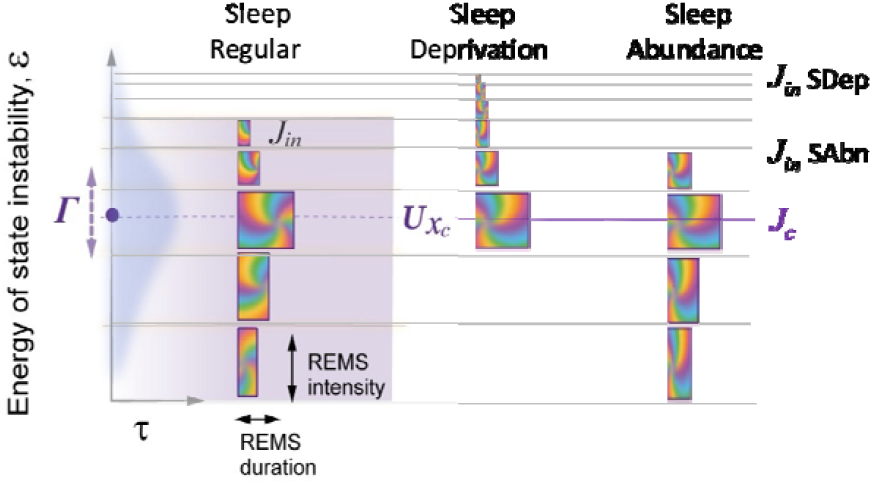
The wave model predictions of the effects of sleep deprivation and sleep abundance on the duration and intensity of REMS episodes. *Regular sleep:* Maximal REMS duration (width of the multicolor blocks) occurs near the resonance level *J*_*c*_ at the homeostatic energy threshold (*Ux*_*c*_, black dot, dashed line). The reduction in this experimental measure above and below the resonance level (gray area) is defined by the width of the resonance (vertical double-arrow). REMS episode intensity (height of the multicolor blocks) follows linear increase in energy gaps between levels (horizontal gray lines) of the Morse potential. *Sleep Deprivation (SDep):* following prolonged Wake state, sleep starts at higher initial energy level (*J*_*in*_ SD), resulting in smaller initial duration and intensity of REMS episodes. If recovery sleep is of regular duration, awakening may occur before reaching or at the homeostatic threshold level, preventing bell-shaped pattern of REMS episode durations. *Sleep abundance (SAbn):* following brief Wake state, with no prior sleep deficit, sleep is initiated at a lower initial energy level (*J*_*in*_ SA), increasing initial duration and intensity of REMS episodes. The recovery sleep can be shorter than usual due to the relaxing energy of instability reaching the homeostatic energy threshold faster than in regular sleep.

Another important prediction of the model for the effects of sleep deprivation is that recovery sleep of regular duration initiated at habitual bedtime may not have enough time to reach below the homeostatic energy threshold and thus the resonance level. As a result, the REMS duration curve would not manifest a bell-shape but would show only continuous increase (Fig. 10, SDep).

In contrast to the effects of sleep deprivation, when sleep onset at habitual bedtime follows prior acute sleep abundance, such as due to a prolonged daytime nap, sleep initiation would be associated with lower energy level, larger and a smaller difference between *j*_*in*_ and (Fig. 10, SAbn). These accurately predict that sleep would start with lower NREMS intensity, higher REMS intensity, and longer REMS episode duration, respectively.

## Discussion

This study introduces a novel model concept of sleep dynamics, based on the interaction of two probability waves that represent Sleep and Wake states. The model is systematically tested against experimental data on normal sleep patterns, demonstrating high statistical significance in predicting the dynamics of four primary sleep measures representing both NREM and REM sleep. Furthermore, the experimental validation of the predicted Sleep Cycle Invariant strengthens the credibility of the model. These findings provide compelling evidence of the intrinsic unity between NREM and REM sleep, a concept for which proof has been sought by many research groups for seven decades since the discovery of REM sleep^5^.

The model utilizes advanced mathematical apparatus of wave mechanics, yet it produces simple analytical formulas for the major sleep measures, enabling efficient analysis of large datasets. This simplicity arises from the direct relationship between the four specific characteristics of the quasi-periodic process, which evolves over consecutive energy levels of the Morse potential, and the four primary sleep measures of each sleep cycle. These include the period of oscillation (NREM duration), its amplitude (NREM intensity), the energy gap between levels (REM intensity), and the time it takes to transit that gap (REM duration). The wave model proves equally effective when applied to independent datasets collected under regular or extended sleep opportunities. Additionally, it accurately predicts the effects of sleep deprivation or abundance on sleep architecture.

The wave model proposes that the sequential occurrence of NREM sleep followed by REM sleep within each sleep cycle reflects their distinct yet complementary roles. During NREM sleep, the time evolution of the wave packet prepares the energy for release. As the system enters the region of equilibrium between Sleep and Wake states, the strong interaction between the two waves leads to their coherent superposition, resulting in REM sleep. This is when the primed fraction of the energy is released, followed by further evolution of the wave packet over the subsequent NREMS episode to prepare more energy for release. Notably, the initial proposal by Eugene Aserinsky, the discoverer of REM sleep^5^, that REM sleep intensity is a measure of “sleep satiety”, finds further support in the wave model, where REM sleep intensity is directly proportional to the energy of instability relaxed within each sleep cycle.

The specific physiological or molecular substrate of what has conventionally been referred to as sleep need, sleep pressure^2^ or state instability^7^ is yet to be fully understood. While a mathematical model cannot provide an answer to this question, an accurate quantitative description of the sleep process can inform and facilitate the exploration of functional links on molecular and systemic levels. This expectation is particularly relevant to the Sleep Cycle Invariant, as the search for invariant relationships has been fundamental to the development of theories and laws in the natural sciences. The discovery of invariants has played a crucial role in advancing scientific knowledge by providing a framework for making critical predictions, designing experiments, and guiding further research.

The representation of REM sleep as a coherent superposition of two waves representing Sleep and Wake states may initially seem unexpected, even though aligns with the accurate quantitative predictions of the model. However, there exists a striking similarity between these two phenomena. In physics, the coherent superposition of interacting waves is known to give rise to a new state that simultaneously exhibits characteristics of both constituent states, rather than simply being a combination or alternation between them^24,42^. This observation is consistent with REM sleep, which manifests a simultaneous presence of features associated with both Sleep and Wake states. Consequently, REM sleep has acquired alternative conventional names such as “paradoxical sleep” or a “third state” of consciousness^43^, highlighting its unique nature.

Throughout successive sleep cycles, the gradual release of energy of state instability brings the system closer to a more stable Wake state. As a result, longer and more intense periods of REM sleep increase the likelihood of spontaneous awakening^31^, as they contribute to a more efficient reduction of state instability. According to the model, the influence of the circadian clock on the probability of transitioning from sleep to wakefulness and the duration of REM sleep episodes^44^, but not on the other three primary measures of sleep architecture, can be attributed to the clock’s modulation of the strength of the Sleep-Wake interaction (Method 1). It is important to note that the present study focuses on investigating typical sleep patterns under regular entrained conditions, while the specific contributions of the circadian and homeostatic forces to the Sleep-Wake interaction will be addressed in a separate report.

The model has only one independent variable, the regulating parameter *x*, a one-dimensional reduction of the overall physiological state that reflects the system’s ability to maintain stability (Fig. 11). The energy of state instability is dependent on the regulating parameter. It increases when the system’s stability is altered, and the regulating parameter deviates from the equilibrium region of the two states, Sleep and Wake (Fig. 2, 11). Similar relationships can be found on many physiological levels, from cellular to systemic. For example, if the heart rate variability (HRV) is one of the indicators of the physiological state stability^13^, some measure of sympathetic nervous system activity, which is known to increase when HRV decreases, could reflect the energy of state instability. The regulating parameter in our model is a simplified version of a combined effect of many such physiological processes that reflect the overall system stability.

**Figure 11.**
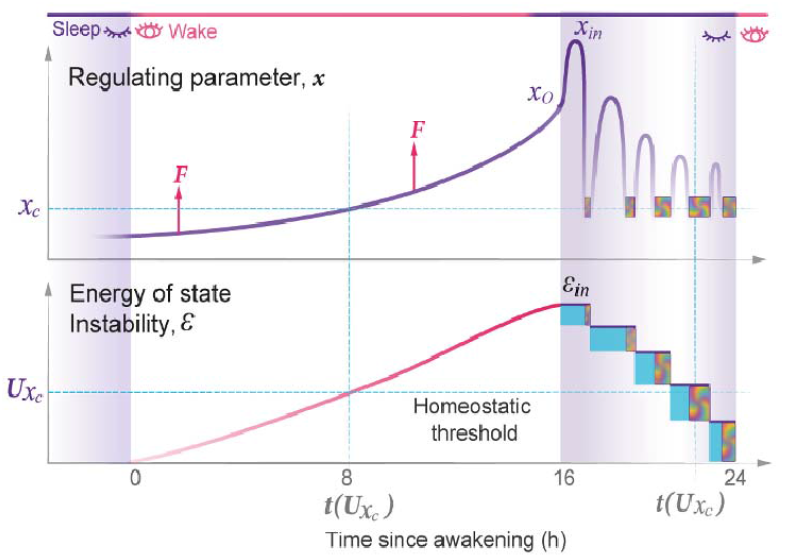
Time-dependent changes in the regulating parameter of state stability and energy of state instability. *Top panel:* Time-dependent changes in the regulating parameter of state stability *x* (vertical axis) over Wake (white area) and Sleep (purple area). In Sleep, the period of *x* oscillations corresponds to the duration of consecutive NREMS episodes, while decline in their amplitude squared to the drop in NREMS intensity. Each cycle ends around *x*_*c*_ (horizontal cyan line), the region of Sleep-Wake equilibrium where REMS episodes occur (multicolor blocks). *F* – the driving force (red arrow) that increases *x*. *Bottom panel:* Time-dependent changes in the energy of state instability (vertical axis) over the Wake state (white area) and during stepwise decline over consecutive sleep cycles (block width - duration: NREMS – cyan; REMS - multicolor). Maximal energy level reached () defines initial NREMS intensity. The linearly increasing portions of energy released correspond to REMS intensity (block height). Horizontal cyan line - potential energy of homeostatic threshold, *U(x*_*c*_). Vertical cyan line – timing of *U(x*_*c*_), with daytime timing arbitrary placed mid-day; horizontal axis – time since awakening (hours).

The homeostatic relationship between the Sleep and Wake states is largely undisputed. However, previous models have not included the homeostatic setpoint or the homeostatic threshold, relying mostly on the limits provided by the circadian clock^2,15^. The homeostatic relationship between Sleep and Wake states is formalized in the wave model of sleep dynamics, which includes a critical homeostatic energy threshold around the equilibrium setpoint of the crossing of the two potential curves. This threshold is the point of equal stability and strong interaction of the Wake and Sleep states and, at the same time, is the site of resonance amplification of the duration of coherent superposition. The parametric position of the homeostatic threshold can be estimated based on the peak of the normal bell-shaped REMS duration curve. Individual differences in this threshold may be significant for understanding variations in normal sleep and sleep disorders.

Finally, we would like to emphasize that the Sleep and Wake probability waves that the model operates with are not of quantum mechanical nature. It is well established that the stochastic classical systems can mimic the dynamics of probability waves observed in quantum physics^25-29^ and we consider our finding to be yet another example of this phenomenon. We propose that the conceptual and mathematical apparatus of wave mechanics can be applied to the study of multiple aspects of both Wake and Sleep states, including changes induced by chronic sleep deprivation, circadian phase shifts, disease, pharmacological or environmental interventions.

## METHODS

In this section, we first outline the analogies between the Sleep-Wake dynamics and the model of a diatomic molecule that informed our work (**a-f**). Next, in Methods **1-15**, we detail the wave model of normal human sleep dynamics and explain how the comparisons between the experimental data and theoretical predictions were made. Method **16** describes the experimental datasets (our and others) that were used to validate the model and includes the Inclusion and Ethics statement for our dataset. Method **17** details the statistical testing.

It is important to note that human sleep is a stochastic process, as reflected in the high inter-individual and night-to-night variation of sleep architecture. Therefore, this work is intended to model average sleep patterns in groups of human subjects studied under well-controlled conditions. Additionally, it should be noted that the wave model in this paper only addresses normal human sleep.

### Analogies between the wave model of sleep and the model of a diatomic molecule

We view the dynamics of the Sleep state as the result of the interaction of two probability waves formed by numerous coherent biochemical oscillators that require coherence. In developing the mathematical approach, we drew a structural analogy with a quantum mechanics model of a diatomic molecule^24^ and relied on several key parallels:

a. *Fast and slow components determine the dynamics of the system*. In a diatomic molecule, changes in electronic states are fast, while changes in the distance between the nuclei of the two atoms (the internuclear distance R) are slow. In our model, changes in biochemical and electrochemical processes forming Sleep or Wake state are relatively fast, while changes in the regulating parameter of state stability *x* are slow and thus analogous to variations in R.
b. *State stability of the fast component*. In the molecular system, the electron component can be in stable (ground) or unstable (excited) state, which are states of different symmetry at a given value of R. Changes in R can result in the swap of symmetry of the state, such that a former ground state becomes unstable, while the former excited state becomes stable. Similarly, in our model, the stability of the Sleep and Wake states is determined by the regulating parameter *x*, and changes depending on variations in *x* value. Low *x* values favor Wake and high *x* values favor Sleep.
c. *The interaction and feedback relationship between the fast and slow components*. For different electronic states in a diatomic molecule, the energy of the fast (electronic) component depends on R, and this dependence creates a potential energy U(R) for the slow nuclear motion. We expect a similar relationship within the Sleep and Wake dynamics, where the parameter regulates the stability of the underlying fast processes, e.g., homeostatic loops. In turn, those fast processes modulate the dynamics of the regulating parameter *x*, creating distinct potential energies *U*_s_(*x*) and *U*_*w*_(*x*) respectively.
d. *Probability waves*. The wave nature of probabilistic processes can be illustrated by the probability waves, or de Broglie waves, which describe the dynamics of the electronic and nuclear components in a diatomic molecule. The non-deterministic nature of the Sleep and Wake processes, as well as the coherent dynamics of their slow and fast components, suggests the use of quantum mechanical analogies in the description of Sleep architecture.
e. *State transitions*. The probabilistic transitions between electronic states of different symmetry, that is the swapping of stable and metastable (excited) states, can occur within certain restricted regions of the R-parameter where the electronic energies of different states have close values. We predict that transitions between the Sleep and Wake states will have similar behavior in the region of crossing or pseudo-crossing of the *U*_*s*_(*x*) and *U*_*w*_(*x*) potential curves. The *x*_*c*_ represents the point of Sleep-Wake homeostatic equilibrium, with U*x*_*c*_ as the homeostatic energy threshold (Fig. 2).
f. *The discrete energy spectra of the stationary probability waves*. In the molecular system, the stationary probability waves have a discrete spectrum of energy for both the electronic and nuclear components, the latter being represented by R-vibrations. In analogy, in our model we introduce the energy parameter *ε* that represents the measure of instability for either Wake or Sleep state. Increase in state instability leads to increase in *ε*.

### Mathematical apparatus of the wave model of sleep dynamics

#### 1. Wave equations for the Sleep and Wake states

It has been demonstrated through both experimental and theoretical studies that classical systems can mimic behaviors commonly associated with quantum mechanics, such as energy level quantization, tunneling, spin structures, and double-slit interference, among others. These effects have been observed in some macroscopic systems containing classical stochastic waves near instability threshold^25-29^. We thus suggested that the probability waves for Sleep (S) and Wake (W) can be also described using a mathematical analogy with probability waves in quantum mechanics. This enabled the use of the two-component Schrödinger equation for the slow nuclear motion in diatomic molecule to describe the motion of the regulating parameter *x* in our model.

The amplitudes and phases of S and W probability waves are described by the time-dependent wave functions *Ψ*_*s*_(*x,t*) * | *S,X* >, and *Ψ*_*w*_(*x,t*) * | *w,x* >, where | *S,X* > and | *w,x* > are the wave functions of the fast intrinsic variables of underlying S and W processes, which are regulated by the *x* variable. These functions, which are analogues of the wave fonctions of different electronic states in a diatomic molecule, are orthogonal at any *x* and *x*’ because of < *x,s*|*w,s’* > = 0. The functions *Ψ*_*s*_(*x,t*) and *Ψ*_*w*_(*x,t*) describe the dynamics of the variable *x* in both S and W States and two-component wave equations can be written in the matrix form:

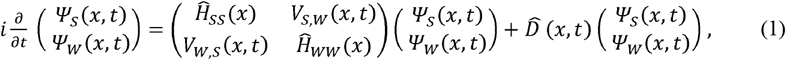

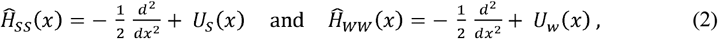

Where *U*_*s*_(*x*) and *U*_*w*_(*x*) are the potential energies modulating propagation of S and W waves, respectively, and 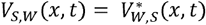 is the matrix element of the operator responsible for the interaction between S and W states. Note that *Vs,w* reflects both the *x*-dependent and time-dependent interstate interaction, including the entrained 24-h periodicity in the Sleep-Wake cycle that is controlled by the circadian system^13^. Efficient S↔W state transitions can be induced by the S-W interaction near the crossing point of the *U*_*s*_(*x*) and *U*_*w*_(*x*) potential curves. Although the potentials *U*_*s*_ and *U*_*w*_ exist simultaneously at the same coordinate *x*, each affects only the corresponding wave. The time, coordinate, and energies in equations 1 and 2 are dimensionless values measured in their specific units. The normalization of the wave functions 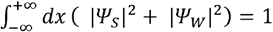 provides the probabilities of realization of S or W state: 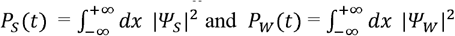.

The operator 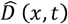 in Eq.1 includes the driving force *F*, which increases the energy of the system, and forces that damp the kinetic energy of the *x*-coordinate. The closest quantum analogy with Eqs.1 and 2 is the system of equations for the nuclear motion of diatomic molecules with two electronic states of different symmetry^24, 45^. In this mathematical analogy, the electronic states represent fast underlying processes regulated by the value of microscopic interatomic distance. However, in the dynamics of macroscopic probability waves^25-29^, as we assume the S and W waves are, the time and spatial scales are many orders of magnitude greater than those described in quantum physics. The time-evolution of wave functions *Ψ*_*s*_(*x,t*) and *Ψ*_*w*_(*x,t*) determines the entire dynamics of S and W states, and consequently the Sleep architecture.

#### 2. S and W stationary waves

Complete sets of eigen functions (stationary waves) can be used as a standard mathematical tool for the determination of wave functions satisfying Equations 1 and 2. The equations for S and W stationary waves can be obtained by neglecting the S-W interaction (*V*_*s,w =*_0) and the action of the 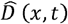 operator. In this case, the stationary S-waves *Ψ*_*s,j*_(*x,t*) = exp(−*I ε*_*j*_ *t*) * *φ* (*x*) and W-waves *Ψ*_*w,k*_(*x,t*) = exp (−*I ε*_*w,k*_ *t*) * *ψ*_*k*_(*x*) are expressed via two sets of independent eigen functions *φ*_*j*_(*x*) and *ψ*_*k*_(*x*), and their eigen energies *ε*_*j*_ and *ε*_*w,k*_, respectively. Since the objective of our model is an accurate description of the Sleep architecture, we mainly consider the stationary S-waves representing the bound states of the potential *U*_*s*_(*x*) with a discrete set of energies *ε*_*j*_ (*j*=0,1,2,3…). The experimental data on the architecture of the W state is limited, and the analysis of W stationary waves is beyond the scope of our model. However, the wave model suggests that the interaction of S and W states plays a key role in the energy relaxation during sleep, specifically in the process of energy release, and that this is reflected in sleep architecture.

#### 3. Energy spectra of the S-waves in the Morse potential

The quasi-periodic nature of the sleep cycles with gradually reducing durations of NREMS episodes *T*_*NR*_ (Fig.1b) indicates that neither parabolic nor rectangular potential wells can describe the Sleep process (Supplemental Fig.2). In contrast, the Morse potential *U*_*s*_(*x*), commonly used in atomic and molecular physics, provides a set of energy levels that are sufficient for accurate quantitative description of the S-wave propagation and the energy relaxation process. The one-dimensional Morse potential is given by the equation:

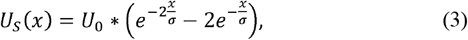

where *U*_0_ and *σ* are positive constants that describe the depth and width of the potential well, respectively. Our model is designed to predict the relative values of *T*_*NR*_ in different seep cycles, so we can scale the depth of *U*_*s*_ potential, *U*_*0*_ = 1. As a result, the energies *ε*_*j*_ at each level and the corresponding eigen wave functions depend on the single parameter σ. The spectra of the discrete energy levels *ε*_*j*_ of the Morse potential are given by the following equation^20^:

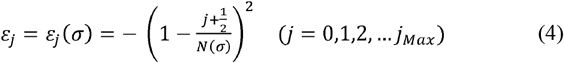

where 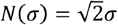 is the dimensionless parameter that regulates the the total number of discrete energy levels in the Morse potential. 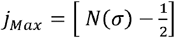.

The discrete energy spectrum and wave functions of stationary waves can also be approximated using quasi-classical Bohr-Sommerfeld model^24,46^. The quasi-classical approximation illustrates the connection between wave and particle dynamics, and describes propagation of the wavepackets of probability waves. We will later consider the dynamics of the quasi-classical wavepacket of S-waves and thus determine the NREMS episode durations, *T*_*NR*._ In the Morse potential, the period of oscillations of the wavepacket moving with an energy close to *ε*_*j*_ declines as the energy *ε*_*j*_ drops, while the energy gaps Δ*ε*(*j*) = *ε*_*j*_ *− ε*_*j − 1*_ between consecutive levels increase (Fig. 2; Supplemental Fig.2).

#### 4. Energy relaxation and structure of the S-wavepacket

Spontaneous W_→_ S transition initiating Sleep generates a wavepacket of S waves with the initial energy 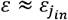. Relaxation of the instability energy *ε* in stepwise transitions, *j* → *j − 1*, occurs through the release of discrete portions of energy Δ*ε*(*j*) = *ε*_*j*_ *− ε*_*j − 1*_. The level *j*(*n*) occupied by the wavepacket during the n-th sleep cycle of the relaxation process is determined by the initial level *j*_*in*_ and the order number *n* of the sleep cycle (Fig. 1, 2), *j*_*in −*_ *n* + 1. In the general case, the exact composition of the wavepacket with an arbitrary energy *ε* includes all stationary S waves belonging to the discrete and continues spectra. At the same time, in the quasi-classical approximation, it can be mostly represented by a single S-wave from *j* energy level, if *ε* is close to the energy level *ε*_*j*_. For simplicity, we neglect the wavepacket dispersion over the entire relaxation process and consider that the wavepacket energy *ε*_*j(n)*_ is reduced only in the stepwise *j*(*n*) → *j*(*n* + 1) transitions during REMS episodes.

#### 5. Quasi-classical motion of the wave packet

At large *j*, the period *T*_*j*_ of quasi-classical oscillations of the center of the wavepacket corresponds to the period of motion of a classical particle with energy *ε*_*j*_ bouncing between the potential walls of *U*_*s*_(*x*). *T*_j_ can be expressed through the energy difference between the neighboring energy levels^46^: 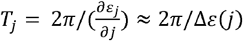, where the derivative 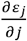 has been replaced with the energy gap Δ*ε*(*j*). The wavepacket motion is also quasi-classical at lower energy levels *j* where the Morse potential is similar to the potential of the harmonic oscillator (equation 3 and Fig. 2). The unitless value of *T*_*j*_ is given by the equation:

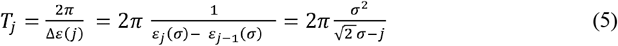

The discrete set of *T*_*j*_ reflects the discrete structure of the energy spectra *ε*_*j*_ of S stationary waves. The set of consecutive *T*_*j(n)*_ periods with *j = j*(*n*) predicts the relative durations of consecutive NREMS episodes *T*_*NR*_(*n*), i.e., periods of the quasi-classical motion in the Morse potential. These values depend only on the width of the potential well *σ* and an actual level *j* occupied by the wavepacket during the *n*-th sleep cycle.

#### 6. Interaction between S and W stationary waves and their coherent superposition

The independence of S and W states is strongly violated within the *δx*_*c*_ region of non-adiabatic behavior around the crossing point *x*_*c*_ (Fig. 2, 5a, 8), where the interaction |*V*_*s,w*_(*x,t*)| is larger than or comparable to the difference between the potential energies: |*U*_*s*_(*x*) − *U*_*w*_(*x*) | ≲ | *V*_*s,w*_(*x,t*) |^23^. Thus, inside 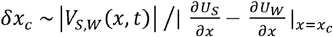, the stationary S and W waves can form a coherent superposition or entanglement. The formation of S-W entanglement inside the moving wavepacket can be clarified by a simplified example in which an entangled state is constructed from isolated stationary S and W waves. A two-state model includes a single S-wave eigenstate *φ*_*s*_(*x*) and single eigenstate of W-wave *ψ*_*w*_ (*x*) with energies *ε*_*s*_ and *ε*_*w*_, and a time-independent potential of S-W interaction *V*_*s,w*_(*x,t*) = *V(x)* can be used to explain this phenomenon. This interaction creates new stationary waves with new energies *E*_*a,b*_ and the wave functions 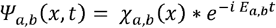, where new eigen functions *χ*_*a,b*_(*x*) are expressed via the coherent superposition of the unperturbed *φ*_*s*_(*x*) and *ψ*_*w*_(*x*) waves of S and W states:

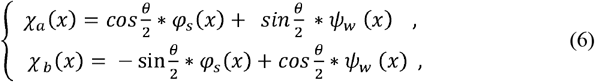

The eigen energies *E*_*a,b*_ and coefficients of the coherent superposition 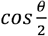 and 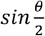 are given by the equations:

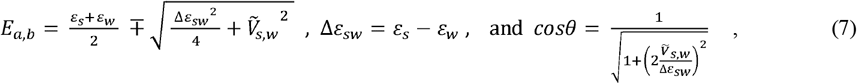

where 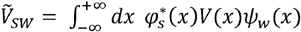 is the matrix element of the S-W interaction potential, and *θ* is the mixing angle. For simplicity, we assumed zero values of the diagonal matrix elements, 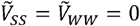. Note that the energies *E*_*a,b*_ of the new states differ from the energy levels of *U*_*s*_ and *U*_*w*_ potentials. The probabilities *p*_*s*_ and *p*_*w*_ to detect the characteristic features of S and W states in the new stationary wave *ψ*_*a*_(*x,t*) are expressed in terms of the coefficients of coherent superposition: 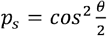 and 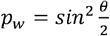. Under condition of weak state interaction, i.e., *θ* ≪ 1, the new stationary wave *χ*_*a*_(*x*) is mainly represented by *φ*_s_ (*x*), occupying primarily the region of S stability *x > x*_c_. The second stationary wave *χ*_*b*_(*x*) is located predominantly within the region of W stability *x* < *x*_*c*_ and it is represented mostly by *ψ*_*w*_(*x*). The parameter of perturbation 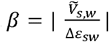 describes the relative strength of the interaction 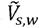 and regulates the fractions of W and S state within their coherent superposition, *ψ*_*a,b*_(*x,t*). In case of *β* → 0, i.e., weak S-W interaction or very large energy difference Δ *ε*_*sw*,_ the entanglement is not formed. In contrast, large value of 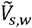 or small difference between S and W energy levels (*β* ≫ 1 or Δ*ε*_*SW*_ → 0) creates conditions for strong coherent mixing of S and W waves. In this case, the mixture angle is 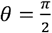, the wave mixture coefficients are 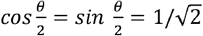, and the coherent S-W superpositions 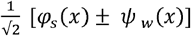 include equal fractions of S and W waves.

The condition of Δ*ε*_*SW*_ → 0 can be seen as a resonance between two stationary waves of different symmetry. Accidental resonance can occur between stationary waves contained within different potential wells^47^. Under conditions of wavepacket propagation, its motion is considered as an adiabatically slow process and the mechanism of formation of coherent states includes similar mixing of the stationary waves, with one important difference: the entanglement of the S and W states is temporary and lasts for as long as the center of the wavepacket remains inside the non-adiabatic region *δx*_*c*_.

#### 7. The coherent superposition of S and W waves and the wavepacket dynamics

In the energy relaxation process, the wavepacket is composed mainly of *φ*_*j*(*n*)_(*x*) waves and its center moves like a classical particle, bouncing between *U*_s_(*x*) walls (Fig. 2). Within this semi-classical approximation, transitions between W and S states can occur when the center of the wavepacket approaches the *δx*_*c*_ region where the crossing of *U*_s_(*x*) and *U*_w_(*x*) potential curves enhances the efficiency of S and W interaction. Within *δx*_*c*,_ the wavepacket can temporarily incorporate a substantial fraction of stationary W waves, as shown in equations 6 and 7. The S-W interaction creates a delay *τ* in the wavepacket propagation, and leads to a new temporary state represented by the coherent superposition of S and W waves. Thus, an organism can simultaneously exist in both S and W states over the time interval *τ*, until the wavepacket leaves *δx*_*c*_ region. In our model, this new state corresponds to REMS, coherently incorporating S and W features. The *τ* defines the lifetime of the coherent superposition within *δx*_*c*_ and thus REMS episode duration *T*_*REM*_.

#### 8. Short time-delay induced in Landau-Zener transition

The presence of the region of the enhanced S-W interaction *V*_*S,W*_, i.e., the region of non-adiabatic behavior of S state, leads to additional accumulation of the phase *η*(*ε*) of the stationary wave with energy ε, and thus the delay in the wavepacket propagation. The resulting time-delay *τ* (*ε*) can be calculated using the energy dependence of the wave phase shift, 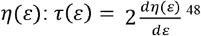. The time delay *τ*_*LZ*_ (*ε*) of the wavepacket passing through *δx*_*c*_ can be also estimated using the semiclassical Landau-Zener model

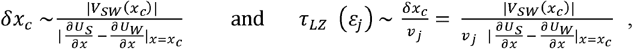

Where *v*_*j*_ is the velocity of the classical particle moving with energy ε*j* through the *x*_*c*_ region. Usually, the Landau-Zener model cannot provide long *τ* because interaction potential V_SW_(*x*) is considered as a small perturbation with respect to *U*_*S*_(*x*) and *U*_*W*_(*x*) In our model, *τ*_*LZ*_ is the background time-delay in the absence of the resonance process. General mathematical analysis and experimental data on the non-adiabatic transitions and phase accumulation for the extended Landau-Zener model have been reported in^49-52^.

#### 9. The resonance enhancement of REMS episode duration and energy release

Within *δx*_*c*,_ the presence of W-wave component of the wavepacket leads to the action of the damping forces included into the operator 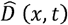 as shown in equation 1. This allows for the wavepacket to release a fraction of its energy, efficiency of the energy Δ*ε*(*j*) corresponding to the energy gap between the *j* and *j*-1 levels. At a constant value of damping forces, the efficiency of the energy release depends on the strength of S-W interaction and the time-delay *τ*(*ε*_*j*_). The value of the released energy Δ*ε*(*j*) increases with the number of cycles *n*. Facilitating energy release requires more time for the action of the damping forces, i.e., increasing *τ*(*ε*_*j*_). This can be accomplished through a resonance condition for the incoming S-wave, if the energy of the resonance level *ε*_*jc*_ is close to U (*x*_*c*_), the potential energies of S and W states near their crossing point *x*_*c*,_ where the strength of S-W interaction is high. The temporary capture of the wavepacket into the resonance state augments the lifetime of coherent S-W superposition and thus T_*REM*_.

#### 10. Mechanisms of resonance formation

The formation of a resonance state can occur through different mechanisms. One example is the Feshbach resonance^53^, in which the energy levels of *U*_S_ (*x*) and *U*_*W*_(*x*) are located near *U x*_*c*_. In this case, *τ* (*τ*_j_) determines the effective time that the W-state is present within the composition of the S-wavepacket when the center of the wavepacket is located within the δ *x*_*c*_ region. Another mechanism by which an S-wave resonance may occur is through a very strong local interaction between the S and W waves, which creates a quasi-bound state localized near *x*_*c*._ In this case, the avoided crossing structure of the adiabatic potentials can support the resonance (quasi-bound) state for incoming S-waves and temporarily localize the wavepacket within δ *x*_*c*._ Detailed mathematical descriptions of these resonance effects have been resonance ε*j*_*c*_ can be formally associated with the resonance level *j*_*c*._

#### 11. Lorentz resonance curve

Independent of the mechanism of the resonance, at any energy ε_*j*_ of the S wavepacket, the resonance time-delay τ(ε_*j*_) is sensitive to the energy difference between ε_*j*_ and the resonance energy ε*j*_*c*._ The time-delay can be calculated from the additional resonance phase *η*(*ε*_*j*_)^45^ with a simple expression for the resonance value of *τ*(*ε*_*j*_) given by the resonance Lorentz formula^48,53^:

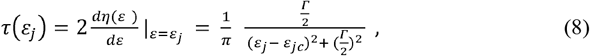

Where ε_*jc*_ and Γ are the energy level and width of the S-wave resonance, respectively. The *j*_*c*_ value can be expressed via 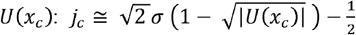 It is not an integer because, in the general case, the resonance energy does not exactly match ε_*j*_ levels. The dynamic changes of *τ(ε*_*j*(*n*)_) over the process of stepwise energy relaxation describe relative durations of consecutive REMS episodes *T*_*REN*_*(n)* over the course of sleep.

#### 12. NREMS and REMS episode durations in absolute time units

The *T*_*NR*_*(n)*,and *T*_*REN*_*(n)* in absolute units (min) can be obtained using equations 5 and 8 for the relative durations *T*_*j(n)*_ and *τ(ε*_*j(n)*_) with *jn=j*_*in*_ ^*_*^ *n +*1,and the absolute time scaling constants A_*NR*_ and A_*REN*_, respectively:

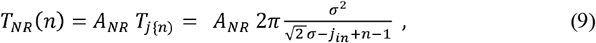

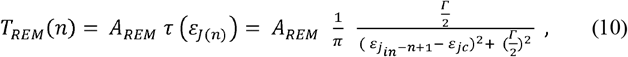

To calculate the relative values of *T*_*NR*_(*n*), two model parameters are needed: the width of *U*_*s*_ potential *σ* and the initial level *j*_*in*_. To calculate the relative values of *T*_*REM*_(*n*), two additional model parameters are required: the energy ε_*jc*_ of the resonance level and the resonance width Γ (Fig. 8). These parameters have been inferred from comparisons of the experimental data on *T*_*NR*_(*n*) and *T*_*REM*_(*n*) and the theoretical predictions of these values given by the analytical formulas in equations 9 and 10 (Supplementary Table 1, Fig. 3, 9). Since the model predicts relative times, the time-scaling constants *A*_*NR*_ and *A*_*REM*_ were used to convert values of relative durations to the absolute time units (min).

#### 13. Predictions of NREMS intensity as a function of the initial energy 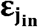 and amplitude of x-oscillations

The NREM intensity *I*_*NR*_(*n*), i.e., the duration of slow-wave sleep (SWS or slow wave activity), is maximal in the first NREMS episode (*n* = 1) and rapidly decreases over consecutive sleep cycles. This behavior correlates positively with the intensity of *x*-oscillations (Fig. 4), presumably because relative intensities of some processes underlying SWS are regulated by *x* and may follow similar oscillatory dynamics. At each level *j* = *j*(n) occupied by the wavepacket in the energy relaxation process, we expect that *I*_*NR*_(*n*) is proportional to the intensity of x-oscillations, i.e., the square of x-amplitudes *L*^2^(*ε*_*J*_): *I*_*NR*_ (*n*) = κ(*ε*_*J*(*n*)_), where the coefficient *κ*(*ε*_*j*(*n*)_) regulates an efficiency of SWS generation over the entire sleep process. The *k*-coefficient should be maximal at the upper *j* level and decline at lower levels: *κ*(ε_*j*_) = const/ ∣ ε_*j*_∣. Note that the energies of the bound waves are negative −1 < ε*j* < 0 and the absolute value ∣*ε*_*j*(*n*)_∣ increases as the level index *j*(*n*) = *j*_*in*_ + 1 − *n* decreases in the energy relaxation process.

The Morse potential is asymmetric and the measure of the amplitude of non-harmonic *<u>x</u>*-oscillations can be a distance *L*(*ε*) between the right and left classical turning points (Fig. 2). For the wavepacket bouncing between the walls of the Morse potential, the oscillation amplitude *L*(*ε*) is expressed through the width of the potential well *σ* and the wavepacket energy ε:

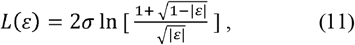

were the value of *ε* in the *n*-th sleep cycle is given by ε_*j*(*n*)_ in equation 4, with *j* = *j*(*n*). Theoretical values of *I*_*NR*_(*n*) can be normalized to the first cycle and results are given by the formula:

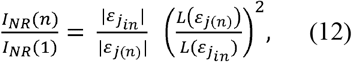

were the wavepacket energies *ε*_*j*(*n*)_ and oscillation amplitudes *L*(*ε*_*j*(*n*)_) are given by equations 4 and 11 with *j* = *j*(*n*). In Figure 4b, theoretical predictions of relative NREMS intensities for different sleep cycles are compared to experimental data obtained from independent groups of healthy volunteers studied under regular sleep conditions and extended sleep conditions (Method 16, below). It is important to note that the theoretical predictions for *I*_*NR*_(*n*) in equation 12 do not include any adjustable parameters, and all necessary model parameters were inferred from the observed durations of REMS and NREMS episodes (Supplementary Table 1, Fig. 3, 9).

#### 14. Dynamics of REMS intensity depends on energy release

According to the model, the rise of REMS *I*_*R*_(*n*) with *n* reflects an increase in the portion of energy Δ *ε*(*j*(*n*)) released within the *n-*th REMS episode, *I*_*R*_(*n*) ∝ Δ *ε*(*j*(*n*)) Theoretical values for *I*_*R*_(*n*) scaled to REM intensity in the first REMS episode (*n* = 1) show linear dependence on the cycle order number *n*:

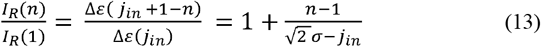

Figure 5b illustrates the theoretical linear dependence of REM intensity on *n*, calculated using equation 13 with the values of *σ* and *j*_*in*_ determined in the analysis of REMS and NREMS episode durations for the regular sleep opportunity group (Supplementary Table 1, Method 16). The slope of the theoretical curve 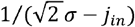 depends on two model parameters through the single combination 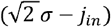 that indicates an approximate number of energy levels above *j*_*in*_. The slope of *I*_*R*_ (*n*) /*I*_*R*_ (1) increases as the initial level *j*_*in*_ approaches the maximal value 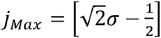 computed without any fitting parameters and are in excellent agreement with experimental data collected by us and others^34,35^ in three independent groups of young healthy subjects, as shown in Figure 5b.

#### 15. Sleep Cycle Invariant

The wave model predicts the existence of a value that remains constant over consecutive sleep cycles, the Sleep Cycle Invariant (SCI). The SCI is defined as the product of the NREMS episode duration, *T*_*j*(*n*),_ and the REMS episode intensity, *I*_*R*_ (*n*):

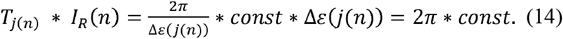

The SCI value does not depend on the value of energy gap, Δ*ε*(*j*) and can serve as a signature of sleep cycle integrity or overall sleep quality. Note that, while calculating SCI for the first sleep cycle, the shorter duration of *T*_*j*(*n*=1)_ (due to *x*_o_ position; Fig. 2) has to be accounted for and the experimental value for *T*_*j*(*n*=1)_ has to be multiplied by 4/3.

#### 16. Datasets

The model was validated against several sets of polysomnographic data collected under well-controlled conditions of sleep laboratory in groups of young healthy subjects with normal sleep patterns. Those presented in Figure 3-5, 7 and 9 are described below:

a. *Regular Sleep Opportunity protocol*. The representative group data for regular sleep shown in Figures 3-5, 7 and 9 were part of our larger study on the circadian regulation of sleep and hormonal functions (“Multimodal Circadian Rhythm Evaluation” PI: IVZ), which will be reported in full elsewhere. The study was conducted in accordance with the Declaration of Helsinki on Ethical Principles for Medical Research Involving Human Subjects, adopted by the General Assembly of the World Medical Association, and approved by the Boston University Institutional Review Board. All the participants provided written informed consent. The subjects were 24 young healthy male volunteers (Mean ±SEM: 24.5 ± 4.4 years of age, ranging19-34 years of age) who were selected based on the following self-reported criteria: 7-9 hours of habitual nighttime sleep, small (<1.5h) changes in sleep length on weekends, no sleep complaints, no history of chronic disorders or regular medications, no recent trans-meridian travel, no drug use, no smoking, habitual coffee consumption not exceeding 3 cups a day. Over the two weeks prior to the inpatient part of the study, the sleep-wake cycle was documented using activity monitors (Phillips Inc.) and a sleep log. Starting on Friday night, subjects spent 3 consecutive nights in the General Clinical Research Center of Boston University School of Medicine. The time in bed was scheduled individually to correspond to the habitual bedtime and subjects were allowed to stay in bed for 9 consecutive hours. Sleep was recorded using polysomnography (Nihon Kohden PSG system), as per standard techniques, and the sleep stages were visually scored for consecutive 30-s epochs^55^. To be included into the regular sleep data set, individual sleep nights had to satisfy the following criteria: sleep efficiency of not less than 85% and the absence of sleep apnea or other symptoms of sleep disorders (n=39 nights total). NREMS-REMS cycles were defined by the succession of a NREMS episode of at least 10 min duration and a REMS episode of at least 3 min duration. No minimum criterion for REMS duration was applied for the completion of the last cycle. A NREMS episode was defined as the time interval between the first two epochs of stage 2 and the first occurrence of REMS within a cycle. A REMS episode was defined as the time interval between two consecutive NREMS episodes or as an interval between the last NREMS episode and the final awakening.
b. *Extended Sleep Opportunity protocol* is described in detail in the original reports by Barbato & Wehr^30^ and Barbato et al.^31^. In brief, the study was conducted in 11 healthy male volunteers, 20-34 years of age. The subjects were studied for 4 weeks, with regular activities over 10 hours of light and bedrest over 14 hours of darkness, when they were encouraged to sleep. The total of 308 sleep records were analyzed. The data used in the present study (Fig. 3-5,7,9) were obtained from Tables 1-3 of ^30^ and Table 2 of ^31^, in consultation with Dr. Barbato.
c. REMS intensity (REM density) data presented in Figure 3e was obtained from the original reports by Aserinsky^34^ (11 normal subjects, young males and females) and Marzano et al.^35^ (50 normal subjects, young males and females). In both studies, the subjects were identified as university students. The REM density data per sleep cycle of baseline night recordings were obtained from p. 550 of ^34^ and Figure 1 of ^35^.

#### 17. Statistical Testing

Goodness of fit was assessed using one-sided *χ*^2^ test, with the degrees of freedom equal to *n* − 1 − *p*, where *n* is the number of independent sleep cycle values being fit, and *p* is the number of model parameters being fit (*p* = 4 for episode duration fits, and *p* = 0 for intensity fits). Note that due to the normalization employed, the number of independent sleep cycle values in each plot is one less than the number of cycles shown. *χ*^2^ and *R*^2^ calculations were carried out using standard R functions

## Author contributions

V.K. and I.V.Z. designed research; I.V.Z performed sleep research; V.K. conducted mathematical modeling; V.K., and I.V.Z. wrote the paper. Authors declare no competing interests.

The dataset generated by the co-author (IVZ) and analyzed during the current study (Method 16, Regular sleep) is available from the corresponding author on reasonable request.

The mathematical algorithms of the wave model of sleep dynamics in “Wolfram Mathematica” format are available from the corresponding author on reasonable request.

## Supplemental materials

**Supplemental Table 1.** The Wave Model of Sleep parameters for groups with Regular and Extended sleep.

| Group | $U_s$<br>width $\sigma$ | Number<br>of initial<br>level $j_{in}$ | Resonance<br>level $j_c$ | Resonance<br>width $\Gamma$ |
| --- | --- | --- | --- | --- |
| Regular sleep | 10 | 10.3 | 7.1 | 0.44 |
| Extended sleep | 11.3 | 11.8 | 7.6 | 0.52 |

Regular sleep: 9-h regular nighttime sleep opportunity, described in Method 16.
Extended sleep: 14-h extended nighttime sleep opportunity, described by Barbato and Wher<sup>30</sup>.

**Supplemental Fig. 1.**
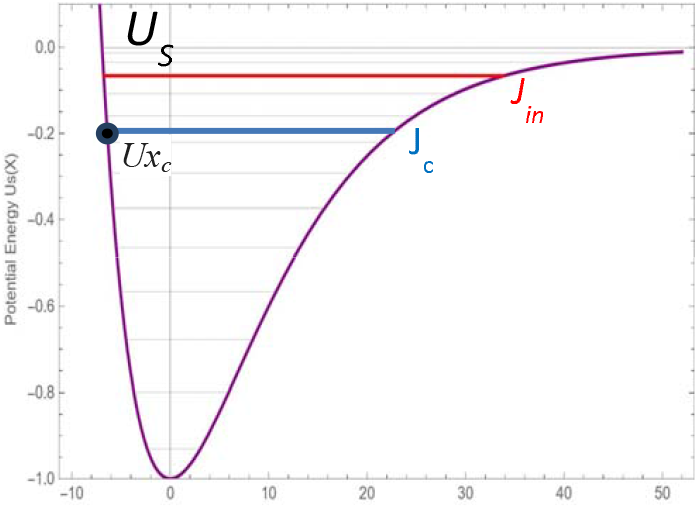
Actual position of the initial and resonance levels within the *Us* potential well. For regular sleep (main Fig. 3b, Extended Table 1), typical position of the initial *Us* energy level (J_in,_ red line) of sleep onset is around level 10. The energy of the resonance level (Jc blue line) is around *Us* levels 7, near the homeostatic energy threshold, *Ux*_*c*_ (black dot). Note that Jc does not belong to *Us* and can be positioned asymmetrically relative to *Us* levels (thin horizontal lines). Conventionally, energy levels are counted from the bottom of the potential well. There is a positive correlation between the total number of *Us* levels and the width of the potential well. In cases of regular sleep, the potential well typically includes around 14 levels.

**Supplemental Fig. 2.**
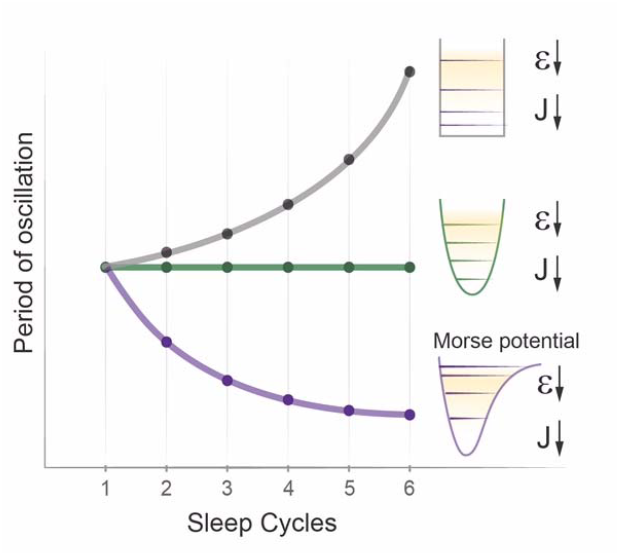
Consecutive periods of energy relaxation in potential wells of different shape. The dynamics of energy ε relaxation over consecutive cycles toward the lowest energy level j depends on the shape of the potential well and energy gaps between levels. The period of oscillation is inverse proportional to the energy gap. For rectangular potential (black), top-to-bottom reduction of energy gaps leads to gradual increase in cycle period during energy decline. For a parabolic well (green), equal energy gaps lead to periods of equal duration (harmonic oscillator). For asymmetric Morse potential (purple), gradual top-to-bottom increase in energy gaps leads to a decline in the period of oscillations during energy relaxation.

